# Cortical polarity ensures its own asymmetric inheritance in the stomatal lineage to pattern the leaf surface

**DOI:** 10.1101/2022.07.15.500234

**Authors:** Andrew Muroyama, Yan Gong, Kensington S. Hartman, Dominique Bergmann

**Author notes:** Co-correspondence to and.

## Abstract

Asymmetric cell divisions (ACDs) specify differential cell fates across kingdoms. In metazoans, preferential inheritance of fate determinants into one daughter cell frequently depends on polarity-cytoskeleton interactions (*1, 2*). Despite the prevalence of ACDs during plant development, evidence for analogous mechanisms that segregate fate determinants during ACD remain elusive. Here, we describe a mechanism in the *Arabidopsis thaliana* leaf epidermis that ensures unequal inheritance of a fate-enforcing polarity domain during the creation of stomata, essential two-celled valves that mediate gas exchange between the plant and environment. Formation of a plasma membrane-associated polarity domain, defined by BREAKING OF ASYMMETRY IN THE STOMATAL LINEAGE (BASL), overrides default division patterns in stomatal precursors. The polarity domain exerts this control by constraining formation of the preprophase band of microtubules that mark the cortical division site and are a hallmark of plant mitosis. Experimentally uncoupling preprophase band establishment from the polarity domain results in aberrant polarity inheritance and subsequent fate errors. Mechanistically, our analyses of the interactions between microtubules and BASL in native and heterologous contexts revealed that the stomatal lineage polarity domain locally depletes cortical microtubules by altering microtubule stability. As the inherited cortical BASL crescent scaffolds a MAPK cascade to suppress progenitor identity in one daughter post-division, we propose that BASL-microtubule interactions represent a novel strategy to link cell identity to division orientation. Together, our data highlight how a common biological module, coupling the cytoskeleton to fate segregation via cell polarity, has been configured to accommodate the unique features of plant development.

## Main Text

Asymmetric divisions prime diverse cell identities by positioning daughter cells relative to fate-enforcing extrinsic signals or by segregating intrinsic fate determinants (*3, 4*). The development of many plant organs depends on ACDs that place daughter cells in proximity to neighbor-derived signals, such as mobile transcription factors (*5, 6*), hormones (*7-9*), or small peptides (*10, 11*). However, whether plants have mechanisms that ensure preferential inheritance of fate regulators remains unresolved.

To interrogate how plant cells couple asymmetric divisions to identity specification, we monitored the cell division and differentiation dynamics of stomatal precursors in the *Arabidopsis* leaf epidermis (Fig. 1A). In this lineage, flexible ACDs in morphologically heterogenous early lineage cells (Fig. 1B) create and pattern stomata by specifying daughters with divergent developmental trajectories. The smaller meristemoid will eventually give rise to the paired guard cells that generate the stomatal pore, and the larger stomatal lineage ground cell (SLGC) will expand to become a pavement cell.

**Figure 1.**
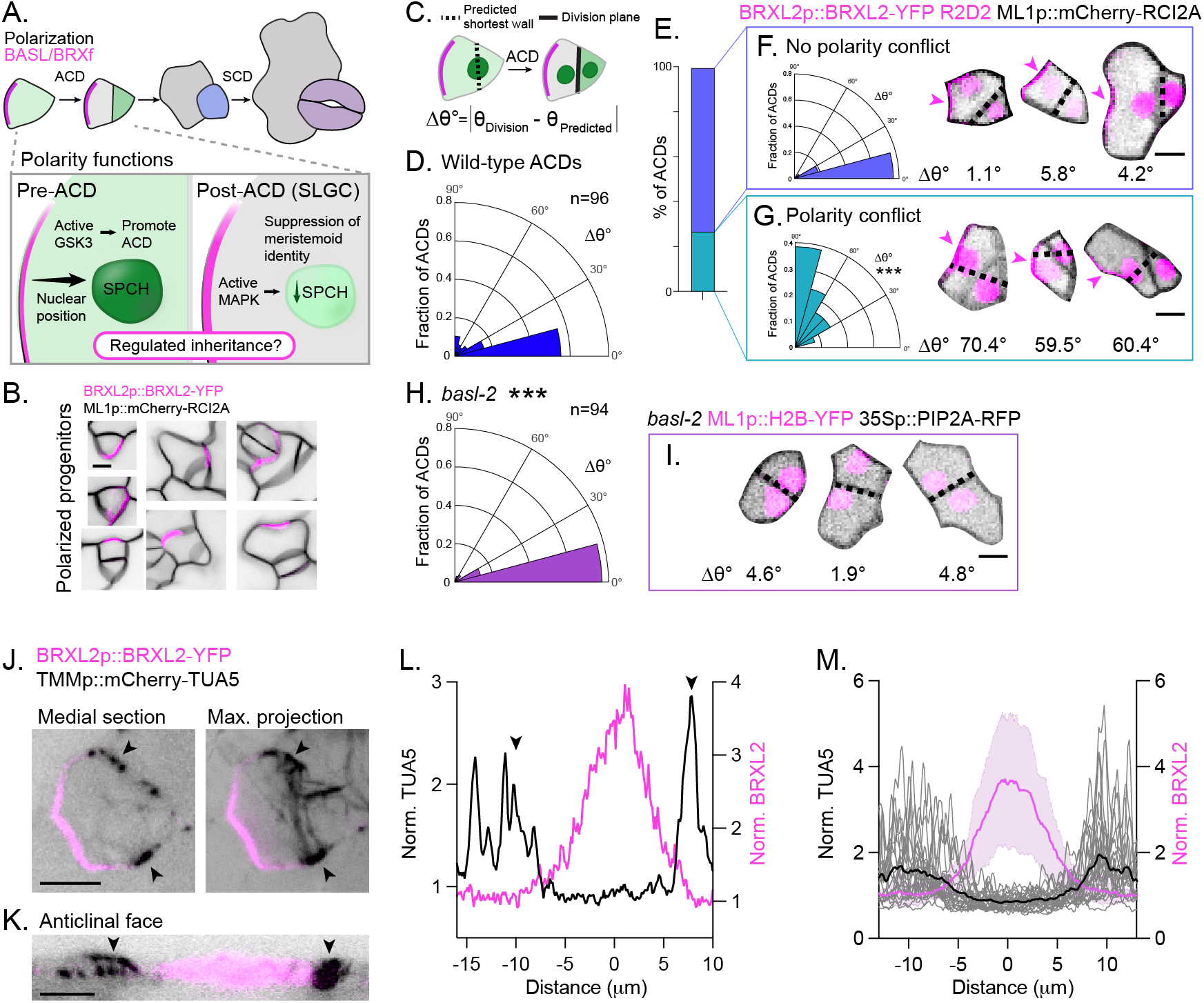
The *Arabidopsis* stomatal lineage polarity domain overrides default division rules during ACD. A. Stomata are created through coupled divisions (ACD and SCD) and fate transitions. (Top) Illustration of the major fate transitions within the *Arabidopsis* stomatal lineage. (Bottom) Illustration of the known functions for the BASL/BRXf domain. Before division (in SPCH+ cells), cortical BASL/BRXf promotes ACD and instructs nuclear migration. Post-division (in SLGCs), inherited BASL/BRXf/YODA phosphorylates MAPK, leading to suppression of the master transcription factor of stomatal identity, SPCH. Whether mechanisms control singular polarity domain inheritance remains unknown. B. Examples of seven polarized stomatal progenitors highlighting the morphological heterogeneity of early lineage cells. Scale bar-5μm. C. The assay used to quantify Δθ°, the angle between the calculated shortest wall and the division plane. This assay was used for the data shown in Fig.1, D to I. For additional details, see Methods. D. Quantification of Δθ° during wild-type ACDs. n = 96 cells. E. The percent of total wild-type ACDs where polarity did (27%) or did not (63%) conflict with the predicted shortest wall. F-G. (Left) Distribution of Δθ° in ACDs where the predicted shortest wall did (G) or did not (F) conflict with the polarity domain. (Right) Three examples of ACDs from their respective classes with associated Δθ°s. The black dotted line marks the calculated shortest wall, and the magenta arrow indicates the polarity domain. Scale bar-5μm. Kolmogorov-Smirnov test comparing the Δθ°s distributions in the two ACD classes: p<0.0001. H. Quantification of Δθ° during progenitor divisions in *basl-2*. Kolmogorov-Smirnov test with WT ACDs: p=0.0005. n = 94 cells. I. Examples of three early-lineage divisions in *basl-2* with associated Δθ°s. The black dotted line marks the predicted shortest wall. Scale bar-5μm. J. Representative images of a medial (left) and maximum projection (right) view of preprophase band (PPB) placement relative to the polarity domain in BRXL2p::BRXL2-YFP TMMp::mCherry-TUA5. Black arrows indicate the PPB. Scale bar-5μm. K. Reslice of the cell in (J) showing microtubule and BRXL2 distribution along the anticlinal wall. Scale bar-5μm. L. Line scan along the cell cortex of the asymmetrically dividing cell shown in (J). The arrows indicate the position of the PPB. M. Microtubule distribution (gray lines) in PPB-forming cells (n=26 cells, average shown in black), aligned to the midpoints of the BRXL2 crescents (magenta line ± s.d.).

All ACDs within the lineage are preceded by the formation of a plasma membrane-associated polarity complex defined by BREAKING OF ASYMMETRY IN THE STOMATAL LINEAGE (BASL) (*12*) and BREVIS RADIX family (BRXf) (*13*) proteins (Fig. 1, A and B). Before mitosis, the BASL/BRXf crescent 1) recruits POLAR LOCALIZATION DURING ASYMMETRIC DIVISION AND REDISTRIBUTION (POLAR) (*14*) to enrich cortical GLYCOGEN SYNTHASE KINASE3 (GSK3) (*15*) and promote ACD, and 2) directs nuclear migration to bias the division site (*16*). After it is inherited by the SLGC through the ACD, cortical BASL/BRXf enhances MITOGEN ACTIVATED PROTEIN KINASE (MAPK) signaling to suppress meristemoid identity (*15, 17*). Despite its central role in coordinating ACD entry and daughter-cell identity post-division (*18*), how BASL/BRXf asymmetric inheritance is regulated is unknown (Fig. 1A).

We performed time-lapse imaging of developing cotyledons harboring markers for nuclei (R2D2 (*19*)), the plasma membrane (ML1p::mCherry-RCI2A) and the polarity crescent (BRXL2p::BRXL2-YFP). In agreement with previous studies, all ACDs resulted in single inheritance of the polarity domain. Close analysis of these cells, however, revealed two ACD subclasses that were defined by the differences in their division planes. The majority (73% of ACDs) divided along the calculated shortest distance that intersected opposing cell walls at the site of the nucleus (small Δθ°, see Methods) (Fig.1, C to F), following the expectations set by the observed division planes in many plant cell types (*20, 21*). For this class of ACDs, polarity-directed nuclear migration (*16*) coupled with minimization of the division plane accurately predicted the final division site.

Division planes in the second class of ACDs (27% of ACDs) deviated significantly from the predicted shortest wall (Fig. 1, D, E and G), suggesting that additional factors impinge on these early lineage divisions. In several *Arabidopsis* tissues, including the shoot apical meristem and early embryo, external stress and hormones can influence cellular geometry to override the shortest wall rule (*22, 23*). The morphological heterogeneity of stomatal precursors was well-represented within both ACD subclasses (Fig. 1, F and G), suggesting that unique geometric features do not define ACD subtypes. Instead, the second class of ACDs shared significant deviations from the calculated shortest division plane (large Δθ°) when that wall was predicted to bisect the cortical polarity site (Fig. 1E). This correlation suggested that polarized BASL/BRXf may be a cell-intrinsic cue capable of overriding default division patterns to control its own asymmetric inheritance.

Next, we tested whether cell polarity is necessary to stratify the two ACD classes by analyzing progenitor divisions in *basl* mutants (*basl-2* 35Sp::PIP2A-RFP ML1p::H2B-YFP). Loss of cellular polarity in *basl* collapsed the two ACD classes into a single one (Fig. 1, H and I) that varied significantly from the total wild-type ACDs (Kolmogorov-Smirnov test, p=0.0005) but not from the first class of wild-type ACDs where there was no conflict between the calculated shortest wall and the polarity site (p=0.1568). Therefore, the polarized BASL/BRXf domain is required to override default division patterns during formative asymmetric divisions.

To determine the basis of this control, we examined cortical microtubule arrays, which play essential roles during division orientation in plant cells (*24*). We found that the preprophase band of microtubules (PPB), which predicts the eventual cortical division site, never formed within the polarity domain (Fig. 1, J to M). An analysis of the Δθ° between the PPB and calculated shortest wall showed a similar bifurcation of ACDs into two classes: the majority (66%) had PPBs that closely aligned to the predicted shortest wall while PPBs in the second class (34%) deviated significantly from the shortest distance (fig. S1). In this second class, 1) PPBs did not form along the shortest wall when the shortest wall bisected the polarity domain, and 2) polarity was required for PPB realignment away from the shortest wall (fig. S1). Together, these data indicated that the BASL/BRXf polarity crescent might orient divisions by controlling PPB placement.

To test this hypothesis, we generated lines to monitor BRXL2 inheritance in the *trm678* mutant (*trm678* BRXL2p::BRXL2-YFP ML1p::mCherry-RCI2A), which does not form PPBs (*25*). In contrast to wild-type ACDs, where BRXL2 was inherited by a single daughter cell, new cell walls frequently bisected the polarity site in *trm678* (32% of ACDs) (Fig. 2, A and B), showing that the PPB is required to ensure complete inheritance of polarized BASL/BRXf. Next, we tracked cell fate outcomes following incorrect BASL/BRXf inheritance by monitoring progression through the stomatal lineage by tracking MUTE (*26*), the initiator of guard mother cell identity, and the subsequent creation of paired guard cells. *trm678* ACDs where BRXL2 was correctly inherited by a single daughter cell showed normal progression; MUTE expression was detectable in the smaller cell after the ACD, and all tracked MUTE^+^ cells in *trm678* became paired guard cells (145/145 MUTE^+^ cells) (fig. S2). In contrast, *trm678* cells where cortical BRXL2 was bisected by the nascent division plane tended to generate daughters that 1) both inherited cortical BRXL2, 2) never transitioned to MUTE^+^ cells, and 3) failed to become pavement cells (Fig. 2C). In agreement with these tracking data, 7dpg *trm678* cotyledons had fewer stomata and a mispatterned epidermis (Fig. 2, D and E). Therefore, the PPB serves as an essential link between the polarity domain and division orientation to regulate stomatal identity.

**Figure 2.**
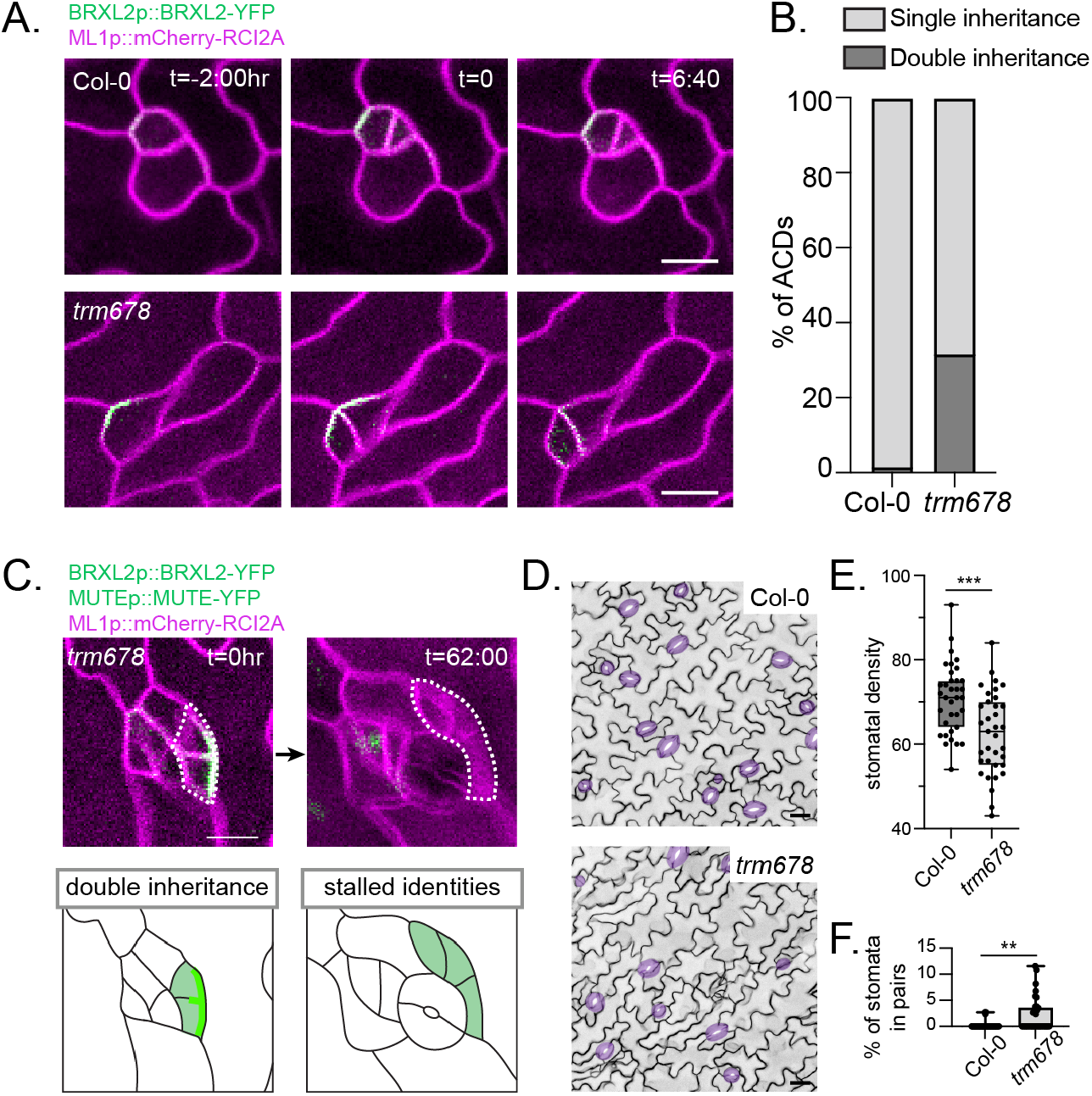
Singular inheritance of the BASL/BRXf polarity domain via PPB placement is required for stomatal patterning. A. Stills from representative time-lapse movies showing division orientation in Col-0 (top) and *trm678* (bottom), relative to the BRXL2-YFP-marked cortical polarity domain. Scale bars-10μm. B. Quantification of Col-0 (n=53) and *trm678* (n=116) ACDs with singular inheritance (BRXL2 inherited by a only one daughter) or double inheritance (BRXL2 inherited by both daughter cells). C. Representative time-course images, with associated cartoons, showing an instance of double inheritance and cell fate stalling 62 hours later from a *trm678* seedling. Scale bar-25μm. D. Representative images of the epidermis of 7dpg Col-0 and *trm678* cotyledons expressing ML1p::mCherry-RCI2A. Stomata are pseudo-colored purple. Scale bars-25μm. E. Quantification of stomatal densities (number of stomata per 581.82μm x 581.82μm area) in 7dpg Col-0 and *trm678* cotyledons (35 seedlings each). p=0.0002. F. Quantification of the percent of paired stomata per 581.82μm x 581.82μm area in 7dpg Col-0 and *trm678* cotyledons (35 seedlings each). p=0.0012.

How does the BASL/BRXf crescent influence PPB establishment? By creating a stomatal lineage-specific microtubule reporter line (TMMp::mCherry-TUA5), we could analyze microtubules and the BRXL2 polarity domain along anticlinal walls with high resolution (Fig. 3A). Unexpectedly, anticlinal microtubules were strongly depleted from the plasma membrane within the polarized domain, even in interphase (Fig. 3, A to C). We confirmed that identical depletions were seen using a BASL reporter and in true leaves (fig. S3). POLAR, which shows overlapping but distinct localization from BASL/BRXf (*14*), co-localizes with microtubules outside the BASL/BRXL2 domain (fig. S3), indicating that microtubule depletion is correlated specifically with BASL/BRXf and is not a generalized activity of polarized proteins in the stomatal lineage.

**Figure 3.**
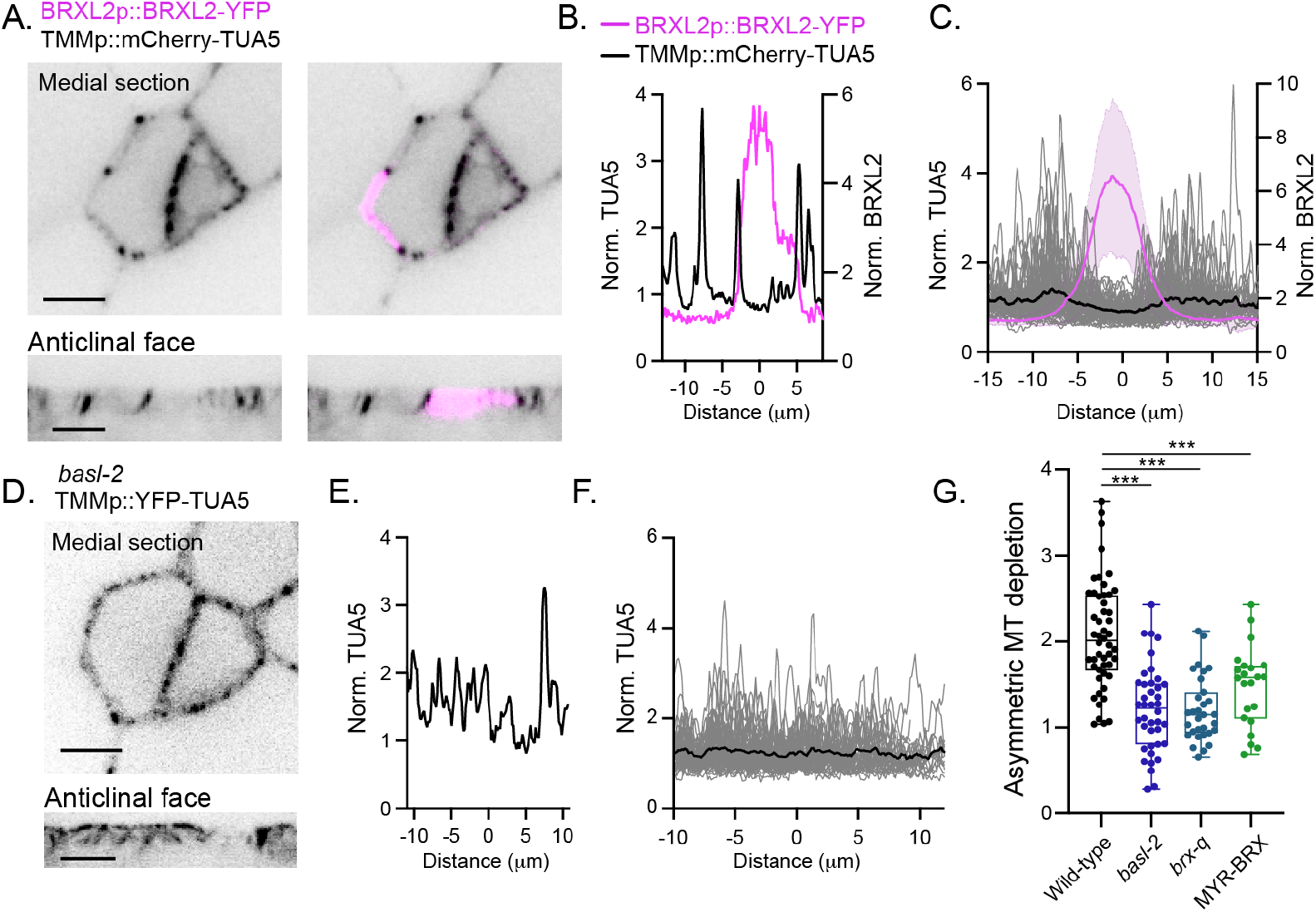
The BASL/BRXL2 domain locally depletes cortical microtubules from the plasma membrane. A. (Top) Representative medial section of a polarized SLGC and associated meristemoid in BRXL2p::BRXL2-YFP TMMp::mCherry-TUA5. (Bottom) Reslice showing microtubule and BRXL2 distribution along the anticlinal face of the same cell. Scale bars-5μm. B. Line scan along the cell periphery of the SLGC shown in (A) showing the normalized BRXL2 and TUA5 fluorescence intensities. C. Microtubule distribution in SLGCs (gray lines) (n=72 cells), aligned to the midpoints of the BRXL2 crescents. The black line is the average of the plotted microtubule signals. D. (Top) Representative medial section of stomatal lineage cells in *basl-2* TMMp::YFP-TUA5. (Bottom) Reslice showing microtubule distribution along the anticlinal face of the same cell. Scale bars-5μm. E. Line scan along the cell cortex of the larger daughter cell in (D) showing the normalized TUA5fluorescence intensity. F. Microtubule distribution in *basl-2* SLGCs (gray lines) (n=58 cells). G. Asymmetric MT depletion scores, calculated from integrated fluorescence intensities in polarized and unpolarized cortical domains (see Methods) in wild-type (n=50), *basl-2* (n=40), *brx-q* (n=31), and BASLp::MYR-BRX (n=22) SLGCs.

BASL/BRXf could 1) locally deplete microtubules or 2) opportunistically polarize to already microtubule-poor regions. To distinguish between these possibilities, we compared microtubule distribution in wild-type and polarity-defective SLGCs. Microtubule distribution was more homogenous in *basl-2 and brx-q (13)*, which abrogate polarity, and in lines where addition of a myristoylation signal (BASLp::MYR-BRX-YFP (*13*)) renders BRX localization largely uniform (Fig. 3, D to G, fig. S4). We also followed unmanipulated SLGCs as they lost polarized BASL/BRXf several hours after ACD. Our time course analysis showed that anticlinal microtubules reappeared within previously polarized regions in mature SLGCs (fig. S4). From these data, we conclude that BASL/BRXf polarity creates and is required to maintain local microtubule loss at the plasma membrane.

Mutual inhibition by opposing plasma membrane-associated domains can drive polarization, as in the conserved PAR networks in animals (*27*) or a recently described polarity system in the monocot *B. distachyon (28)*. Because our data raised the possibility that microtubules and cortical BASL/BRXf could operate in an analogous manner and inhibit the spread of each other, we next tested whether altering microtubule distribution affected the stomatal lineage polarity domain. In agreement with previous results (*29, 30*), we found that microtubules are not necessary for the formation of a polarized BASL/BRXf domain (fig. S5). However, quantification revealed a slight but significant spread of the polarity domain along the anticlinal wall in the absence of microtubules (fig. S6). Short plasmolysis treatments, which dramatically disrupt cortical microtubule distribution, similarly altered polarity boundaries and polarity domain size without complete depolarization (fig. S5). Therefore, microtubules shape BASL/BRXf boundaries in a manner reminiscent of ROP GTPase domain sculpting by microtubule arrays in other plant tissues (*31-33*).

How do cortical BASL/BRXf locally deplete cortical microtubules? Owing to the technical challenges associated with monitoring dynamic microtubule behavior along the anticlinal wall of meristemoids, we created a heterologous system where we could track microtubule dynamics co-incident with the BASL polarity domain. By introgressing a ubiquitous microtubule reporter (35Sp::mCherry-TUA5) into a line expressing a hyperactive version of BASL capable of rescuing the *basl* phenotype (35Sp::GFP-BASL-IC, hereafter referred to as BASL^ectopic^) (*12*), we could monitor microtubule organization relative to BASL in the hypocotyl epidermis. BASL^ectopic^ locally depleted microtubules along anticlinal walls in the hypocotyl epidermis as in the stomatal lineage (fig. S6), demonstrating that this ectopic system recapitulates the molecular interactions found in the leaf epidermis.

BASL^ectopic^ domains extended onto the apical surfaces of hypocotyl epidermal cells and locally depleted cortical microtubules (Fig. 4, A and C), allowing us to observe the BASL-mediated effects on microtubules with precision not possible within the stomatal lineage. Increased microtubule severing has been identified as a key reorganizer of cortical microtubule arrays during several developmental transitions (*34, 35*). However, as severing preferentially occurs at microtubule crossover sites (*36, 37*) and there were few crossovers in microtubule-depleted BASL^ectopic^ regions, severing was largely suppressed within BASL^ectopic^ domains (fig. S7). Tracking of microtubule minus ends within BASL^ectopic^ also indicated that local microtubule depletion was not due to decreased minus end stability (fig. S7). Instead, we found that BASL^ectopic^ had two effects on microtubule plus-ends. First, plus-end polymerization and depolymerization rates were significantly suppressed within BASL^ectopic^ (Fig. 4, E and F, fig. S7). Second, we observed that microtubule plus-ends rapidly underwent catastrophe upon entering the BASL^ectopic^ domain (Fig. 4, B and D). Increased catastrophe rates often lead to complete loss of the microtubule, reestablishing the microtubule depletion zone.

**Figure 4.**
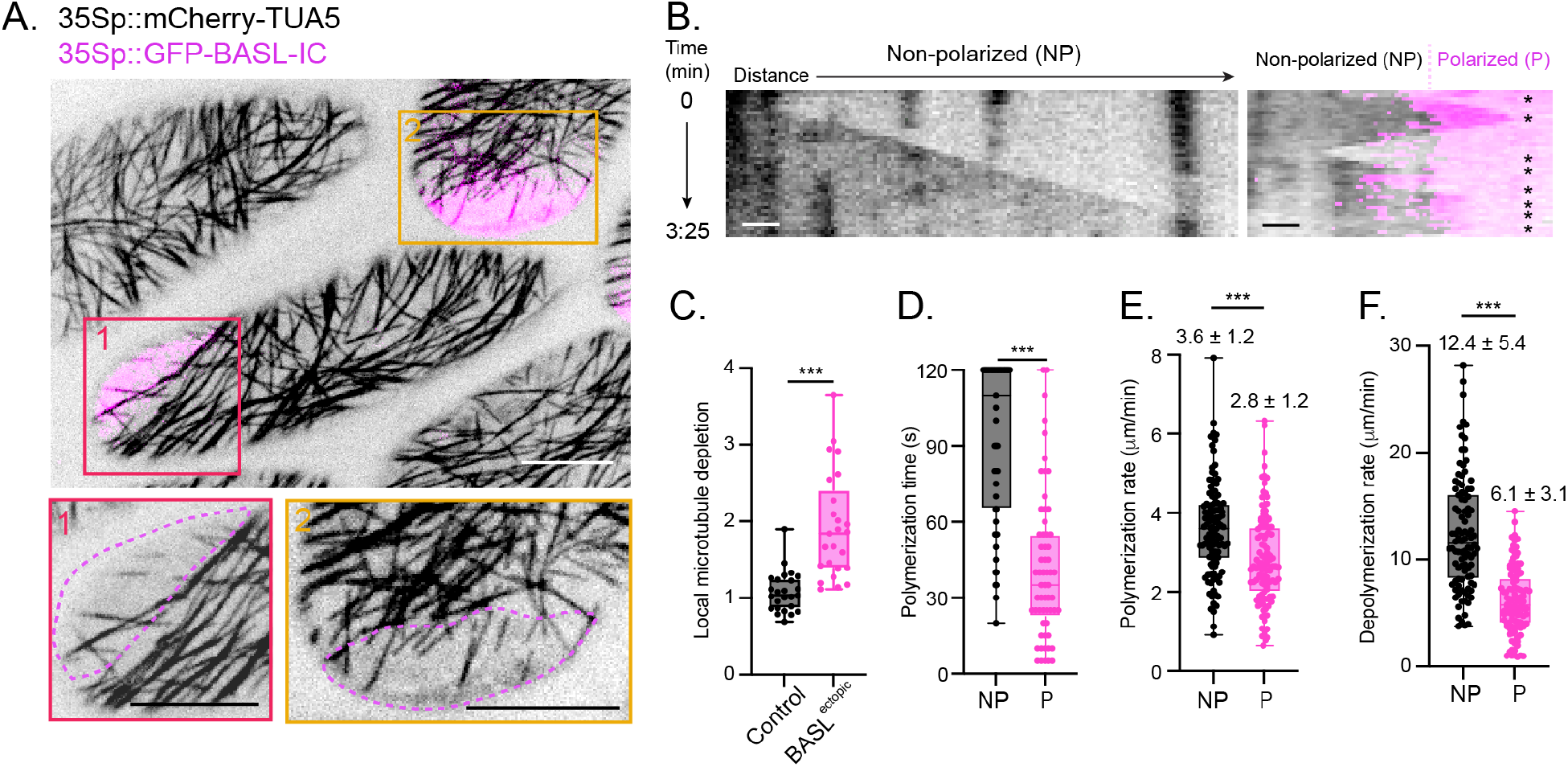
Polarized BASL destabilizes microtubule plus-ends to locally bias array organization. A. Representative images of the apical surfaces in epidermal cells of 35Sp::GFP-BASL-IC 35Sp::mCherry-TUA5 hypocotyls. The boxed regions below highlight the microtubule organization within the polarized domains of two cells. Scale bars-10μm. B. Kymographs showing microtubule plus-end dynamics within a non-polarized region (left) and an ectopic polarity domain (right). Asterisks indicate microtubule catastrophes and rescues. Scale bar-1μm. C. Local microtubule depletion within apical BASL^ectopic^ domains and comparably sized random regions in control hypocotyls. The local microtubule depletion was derived using 35Sp::mCherry-TUA5 fluorescence (see Methods). n=25 hypocotyls for each. D-F. Quantification of microtubule plus-end dynamics within non-polarized (NP) and polarized (P) apical domains. Growth persistence (D), polymerization rate (E) and depolymerization rate (F) were quantified.

To validate that our BASL^ectopic^ findings in the hypocotyl reflect BASL-microtubule interactions within the stomatal lineage, we used two independent approaches. First, we performed time-lapse imaging of stomatal progenitors in BRXL2p::BRXL2-YFP TMMp::mCherry-TUA5 seedlings and observed transient microtubules within the polarity domain that were present only at single time points (fig. S8). Second, to monitor growing plus ends with higher precision along the anticlinal wall, we generated a stomatal lineage-specific END BINDING PROTEIN 1b (EB1b) reporter (TMMp::EB1b-mCherry) and introgressed it into the BRXL2 reporter line. Fewer EB1b puncta were observed within the native polarity domain in SLGCs than in non-polarized regions of the same cells (fig. S8). Therefore, our analyses of microtubule dynamics in native and heterologous systems reveal that the BASL/BRXf domain destabilizes microtubule plus ends to locally deplete them from the polarized region.

As encoders of spatial information, polarity domains are central regulators of ACD in diverse plant tissues (*38-41*). Here, we provide evidence that the BASL/BRXf polarity domain robustly serves dual functions to orient ACDs and specify cell identity by controlling its own inheritance via negative interactions with the microtubule cytoskeleton. Polarity-microtubule interactions now emerge as a common theme to guide asymmetric inheritance of fate regulators during both metazoan and plant ACD, albeit through fundamentally different mechanisms (fig. S9). In the canonical ACD pathway in animal cells, the cortical polarity domain is responsible for 1) localizing fate determinants to one pole and 2) subsequently directing the division angle to ensure their singular and asymmetric inheritance by exerting pulling forces on astral microtubules (*42, 43*). The model we advance here differs in several significant ways. First, the proposed mechanism utilizes core plant-specific mitotic structures without the need to invoke a role for astral microtubules, the presence of which remains unclear during plant mitosis. Second, rather than ensuring its singular inheritance by precisely specifying the ultimate division plane, BASL/BRXf renders a region of the membrane unavailable as the division site. Third, while this mechanism has an identical outcome—asymmetric inheritance of key fate regulator—it is uniquely suited for a morphologically heterogenous population that, nonetheless, must robustly couple ACD orientation with subsequent daughter cell identities.

Of all the documented, polarity-mediated ACDs in plants, those that violate the shortest wall rule, such as those in the early *Arabidopsis* embryo or subsidiary mother cell divisions in *Zea mays*, may be the closest corollaries to the system presented here. While BASL is both eudicot (*44*) and stomatal lineage specific, BRX family proteins participate in additional cellular decisions in Arabidopsis and are much more deeply conserved in the green lineage (*45*), hinting that other tissue-specific regulators could provide context specificity to a common polarity core. Further analysis of polarity will help clarify whether this mechanism is shared across plant tissues and species or whether it has evolved for the particular challenges associated with flexible patterning in the eudicot stomatal lineage.

## Acknowledgments

We thank Dr. David Bouchez for kindly sharing the *trm678* seeds and Amy Lanctot for the POLARp::POLAR-YFP seeds. We thank Dr. Nidhi Sharma for technical help associated with generating the *trm678* reporter lines. We thank Dr. Eric Griffis at the Nikon Imaging Center at the University of California, San Diego for assistance with several imaging experiments. We thank Dr. Mark Estelle, Dr. Alexandra Dickinson, and members of the Bergmann lab for feedback on the manuscript.

## Author Contributions

A.M. and D.C.B. conceived of the study and designed the experiments. Y.G. acquired the time-lapse data for cell tracking in *trm678*. K.S.H acquired and analyzed the *trm678* mutant phenotype data. A.M. performed all other experiments and analyses. A.M. and D.C.B wrote the manuscript with feedback from Y.G. and K.S.H.

## Methods

### Plant material and growth

*Arabidopsis thaliana* seeds were sterilized in 20% bleach with 0.1% Tween-20 for 10 minutes, washed three times with dH_2_O, and plated onto ½ MS (Murashige Skoog) plates with 0.5% sucrose. Plates were stratified in the dark for at least two days at 4°C. Seedlings were grown on plates under long day conditions (16hr light/8hr dark) at 22°C.

All *Arabidopsis thaliana* lines used in this study were in the Col-0 background. Previously reported *Arabidopsis* lines used in this study include: R2D2 BRXL2p::BRXL2-YFP ML1p::mCherry-RCI2A (*1*), *basl-2* ML1p::H2B-YFP 35Sp::PIP2A-RFP (*1*), *basl-2* (WiscDsLox264F02) (*2*), brx*-q (3)*, BASLp::MYR-BRX-YFP (*3*), BRXL2p::BRXL2-YFP (*3*), 35Sp::GFP-BASL-IC (*2*), 35Sp::mCherry-TUA5(*4*), BASLp::YFP-BASL TMMp::mCherry-TUA5 (*1*), *trm678 (5)*, BRXL2p::BRXL2-YFP 35Sp::mCherry-TUA5 (*6*).

BRXL2p::BRXL2-YFP TMMp::mCherry-TUA5, BASLp::MYR-BRX-YFP TMMp::mCherry-TUA5, and POLARp::POLAR-YFP TMMp::mCherry-TUA5 were generated by introducing TMMp::mCherry-TUA5 R4pGWB601 (*1*) into the respective parental lines via agrobacterium-mediated transformation. T1s were selected on ½ MS plates containing 50mM phosphinothricin. To generate *basl-2* TMMp::YFP-TUA5 and *brx-q* TMMp::YFP-TUA5, TMMp::YFP-TUA5 R4pGWB501 was introduced into the parental lines using agrobacterium-mediated transformation. T1 seeds were selected on ½ MS plates with 15mg/mL hygromycin. BRXL2p::BRXL2-YFP (*3*), ML1p::mCherry-RCI2A (*7*), TMMp::mCherry-TUA5 R4pGWB601 (*1*) and MUTEp::MUTE-YFP (*8*) were individually introduced into *trm678* by agrobacterium-mediated transformation. Combinatorial reporters in the *trm678* background were generated by crossing. 35Sp::GFP-BASL-IC 35Sp::mCherry-TUA5 plants were generated by crossing.

### Cloning

TMMp::YFP-TUA5 R4pGWB501 was generated by Gateway cloning (Invitrogen). The YFP sequence was amplified with the following primers: YFP F-5’-gcggccgcatggtgagcaagggcgaggag-3’ and YFP R -5’-gcggccgccttgtacagctcgtccatgc-3’ and subcloned into TUA5 pENTR D-TOPO (*1*) using NotI sites to create YFP-TUA5 pENTR D-TOPO. The TMM promoter in pDONR P4-P1R (540 bp upstream of *TMM*), YFP-TUA5 pENTR D-TOPO, and R4pGWB501 (*9*) were recombined to create TMMp::YFP-TUA5 R4pGWB501.

### Drug and plasmolysis treatment

For oryzalin experiments, 3dpg BRXL2p::BRXL2-YFP TMMp::mCherry-TUA5 seedlings were incubated in liquid ½ MS + 0.75% sucrose with 20mM oryzalin or DMSO for 2 hours. After the incubation, seedlings were mounted for imaging in their respective solutions. For plasmolysis experiments, BRXL2p::BRXL2-YFP TMMp::mCherry-TUA5 seedlings were briefly treated (<10min) in 0.8M mannitol before mounting for imaging.

### Image acquisition

Four microscope setups were utilized for these studies. For time-lapse experiments of division orientation in wild-type (Fig. 1 D to G), *basl-2* (Fig. 1, H and I), and *trm678* (Fig. 2, A to C) and the analysis of fate outcomes in *trm678* (fig. S2), we mounted seedlings in ½ MS + 0.75% sucrose in a custom-fabricated imaging chamber (*10*). The chamber was connected to a peristaltic pump set to a flow rate of 2 mL/hr. Time-lapse movies were acquired on a Leica SP5 with a 25x 0.95 NA dry and 40x 1.1 NA water immersion objectives and HyD detectors using LAS X software. For wild-type and *basl-2* time-lapse movies, images were acquired every 30 minutes. For *trm678* time-lapse movies, images were acquired every 40 minutes.

Phenotypic analysis of the *trm678* mutants was performed using a Leica Stellaris 5 setup with HyD S detectors and a PL APO 20x 0.75 NA objective using LAS X software.

For the analyses of microtubule/PPB distribution (Fig. 1, J to K, Fig. 3, fig. S1, fig. S3, fig. S4, fig. S8), microtubule distribution on the apical surfaces of BASL^ectopic^ (Fig. 4, A and C), oryzalin treatments (fig. S5), and plasmolysis experiments (fig. S5), a spinning disk confocal microscope with a DMI6000 (Leica) stand, Evolve EMCCD (Photometrics) camera, and 100x 1.4 NA objective were used.

For the analysis of microtubule behavior in BASL^ectopic^ cells (Fig. 4B and D to F, fig. S6 and fig. S7), and EB1b dynamics (fig. S8), images were acquired using a SR HP APO TIRF 100x 1.49 NA objective on a Nikon Eclipse Ti2-E microscope with Prime 95B sCMOS (Photometrics) camera.

### Quantification of predicted division angles during asymmetric division (related to Fig. 1, D to I and fig. S1)

To calculate the shortest wall that could be created during asymmetric divisions, xy coordinates defining the cellular outlines and the nuclear centroids were extracted in FIJI from the frame immediately before cell division. Coordinates were imported into MATLAB (version 2021b), plotted, and used to calculate the minimal distance that bisects the cell boundary through the nuclear centroid using distances extracted from calculated polar plots. The calculated shortest wall was visualized on the cell boundary and each cell was individually inspected and validated. The division plane was then identified and measured in FIJI from the frame immediately after cytokinesis. Δθ° is, therefore, the angle between the actual division plane and the calculated shortest wall. An identical approach was used to calculated Δθ° between the PPB and predicted shortest wall.

### Quantification of asymmetric microtubule depletion (related to Fig. 3G)

To calculate microtubule distribution in wild-type SLGCs, fluorescence intensities for both BRXL2p::BRXL2-YFP and TMMp::mCherry-TUA5 were measured along the cell cortex. The area under the normalized microtubule fluorescence signal was integrated in MATLAB and the asymmetric microtubule depletion was calculated by dividing the integrated intensity of the periphery outside the BRXL2 signal by the integrated intensity within the BRXL2 domain. To calculate the microtubule depletion for *basl-2, brx-q*, and BASLp::MYR-BRX-YFP, microtubule intensity profiles were created in FIJI. For each profile, a random region of the cortex ∼10mm in length (comparable in size to the average BRXL2 domain in wild-type cells) was compared to the rest of the cortex.

### Quantification of microtubule dynamics (related to Fig. 4, D to F)

To quantify microtubule polymerization time (Fig. 4D), 2-minute-long time-lapse movies (5 sec. intervals) were used. Only microtubules where the plus-end could be identified for the length of the movie were used for quantifications. To calculate microtubule polymerization and depolymerization rates (Fig. 4, E and F), kymographs were manually generated and used to identify catastrophes and rescues.

### Quantification of local microtubule depletion in BASL^ectopic^ (related to Fig. 4C)

To calculate microtubule occupancy within control (35Sp::mCherry-TUA5) and BASL^ectopic^ (35Sp::mCherry-TUA5 35Sp::GFP-BASL-IC), single optical sections along the apical surfaces of hypocotyl epidermal cells were used. For BASL^ectopic^, TUA5 fluorescence within and outside the BASL domain was calculated and normalized to region area. To calculate “Local microtubule depletion,” the normalized microtubule intensity outside the polar domain was divided by the normalized microtubule intensity within the domain. To calculate a Local microtubule depletion score for control cells, a random region of the apical surface of comparable size to the BASL^ectopic^ domains was created and calculations were carried out in the same manner.

### Quantification of EB1b density (related to fig. S8)

To calculate EB1b density, one-minute-long time-lapse movies of BRXL2p::BRXL2-YFP TMMp::EB1b-mCherry were acquired at 5 second intervals. Line scans of cortical BRXL2 and EB1b intensities were extracted in FIJI, and EB1b puncta were identified with a custom FIJI plugin using the “Find Maxima” function on kymographs. EB1b density was calculated as the number of EB1b puncta per 10mm along the cell cortex per minute.

### Software and statistics

MATLAB (version 2021b) was utilized to generate the polar histograms in Figure 1 and fig. S1. All other graphs were generated in GraphPad Prism 9. All statistical analyses were performed using GraphPad Prism 9. To compare division orientation and PPB angle distributions, the Kolmogorov-Smirnov test was used. For comparisons between two conditions, unpaired t-tests were used except for the EB1b density analysis, where a paired t-test was used. For the analysis of three of more conditions, a one-way ANOVA with Tukey’s multiple comparisons was used.

